# A Computational Model of Mechanical Stretching of Cultured Cells on a Flexible Membrane

**DOI:** 10.1101/2024.06.06.597769

**Authors:** Miles W. Massidda, David Ashirov, Andrei Demkov, Aidan Sices, Aaron B. Baker

## Abstract

Mechanical forces applied to cells are known to regulate a wide variety of biological processes. Recent studies have supported that mechanical forces can cause nuclear deformation, leading to significant alterations in the gene expression and chromatin landscape of the cell. While the stresses and strains applied to cells is it is often known or controlled experimentally on a macroscopic length scale, it is often unclear what the actual forces and displacements are at the microscopic level of the cell. In this work, we created a model of cell deformation during application of mechanical stretch to cultured cells growth on a flexible membrane. This configuration is commonly used is in experimental studies as a means to apply controlled mechanical strains to adherent cultured cells. The parameters used in the study were used for application of strain to a mesenchymal stem cell stretched on a membrane. computational model was created to simulate the stresses and strains within the cell under a variety of stain amplitudes, waveforms and frequencies of mechanical loading with the range of commonly used experimental systems. The results demonstrate the connection between mechanical loading parameters applied through the flexible membrane and the resulting stresses and strains within the cell and nucleus. Using a viscoelastic model of chromatin, we connected the results provide to a rough model of resulting deformation within chromatin from the forces applied to the nucleus. Overall, the model is useful in providing insight between experimentally applied mechanical forces and the actual forces within the cell to better interpret the results of experimental studies.

**Statement of Significance:** In this work, we created a computational model of the mechanical stretching of cell on a flexible membrane under cyclic mechanical loading. This model provides insight into the forces and displacements inside of cell that result from that application of stretch. As many experiments use this set up, our work is relevant to interpreting many studies that use mechanical stretch to stimulate mechanotransduction.

## Introduction

The mechanical environment of a cell plays a crucial role in its functions and behaviors, including cell growth, differentiation, migration, and gene expression.^1–3^ The interplay between mechanical forces and chromatin remodeling has recently emerged as a crucial area of study in cell biology. Mechanical forces not only influence cell shape and movement but also affect chromatin structure and gene expression.^4^ For instance, studies have suggested that external mechanical forces can lead to chromatin remodeling, thereby influencing gene expression.^5^ Conversely, chromatin remodeling can also modulate the cell and chromatin mechanical properties, illustrating the bidirectional nature of this relationship.^6^ In addition, the mechanical environment of the cell can potentially contribute to disease progression by affecting chromatin remodeling.^7^ The role of mechanical forces in influencing chromatin remodeling is also being explored in the context of stem cell differentiation,^8^ cancer,^9^ and aging.^10^ Altogether, these studies and many others highlight the intricate and profound influence of mechanical forces on chromatin structure and function, providing novel insights into cellular processes.

Nuclear mechanotransduction has been found to be an important mechanism in mechanosensing and adaptation of cells.^11^ When a cell is subjected to mechanical force, these forces can be transmitted to the nucleus, leading to deformation of the nucleus and its components. Nuclei are thought to sense and resist the extracellular mechanical stress, protecting the DNA from stress-induced damage.^12^ Recent studies have shown that this deformation may have a significant role in mechanotransduction, leading to extensive changes in the chromatin landscape that causes altered gene expression.^13–15^ In addition, several studies have demonstrated the critical role of physical nuclear deformations in the modulation of cell function,^16^ histone modifications,^17^ transcriptional activity,^18^ and chromatin organization.^19^

Studies on mechanotransduction often apply mechanical forces to cultured cells by stretching the cells on a flexible membrane. This arrangement facilitates imaging of the cells and the use of standard cell culture assays including fluorescent plate reading and western blotting. In addition, it is a practical method for conditioning cells for use in therapeutics and models of disease.^20^ It is, however, less clear how mechanical stretch of the flexible substrate leads to forces on the nucleus. In recent studies from our laboratory, we have found that physiological waveforms of stretch can lead to differential responsiveness of many cell types,^21,22^ and in mesenchymal stem cells (MSCs), lead to increased vascular phenotypes and regenerative properties.^23^ In the these studies, we demonstrated that a very specific types of mechanical stretch induce the MSCs into the vascular phenotypes.

In this study, we created a model of the application of mechanical stretch to a cell on a flexible membrane. This model aims to examine the effects of mechanical stretch of the culture membrane on the actual stresses and deformations that occur within the cultured cell. This study provides a computational model for understanding the forces that different regions of the cells are experiencing when they are stretched at various levels of culture membrane strain. As our previous work has highlighted the importance of physiological waveforms,^21–23^ we examined the differences between the standard sine waveform and a brachial waveform of mechanical stretch that mimics the stretch of the brachial artery during the cardiac cycle. Our previous study demonstrated that brachial waveforms delivered at 0.1 Hz frequency of loading and 7.5%-12.5% maximal strain resulted in MSCs with enhanced vascular regenerative properties and maximal activation of Yap/Taz (Yes-associated protein/transcriptional coactivator with PDZ-binding motif) signaling.^23^ Thus, we examined the effects of changes in frequency and maximal strain as a way to better understand how these types of loading conditions could differentially effect MSC gene expression and mechanotransduction.

## Methods

### Model Parametrization

The computational model was of a single mesenchymal stem cell (MSC) grown on a flexible silicone membrane substrate. MSC mechanical and geometric parameters reported in recent studies were used to parametrize the cell.^24–26^ The cell geometry is represented by an ellipsoid with an elliptical attachment to the surface corresponding to the cell membrane. The nucleus geometry is represented as a smaller ellipsoid embedded within the cell **(SI Fig.1A, B)**. The model defined three main mechanical compartments of the cell: (1) The cell membrane with elasticity of 2.98 +/- 0.04 kPa and viscosity of 4.01 +/- 0.06 kPa*s; (2) The cytoplasm with elasticity of 4.28 +/- 0.33 kPa and viscosity of 5.43 +/- 0.55 kPa*s; and (3) The nucleus with elasticity of 2.01 +/- 0.10 kPa and viscosity of 0.97 +/- 0.08 kPa*s (**Figure 1**).^27^

**Figure 1.**
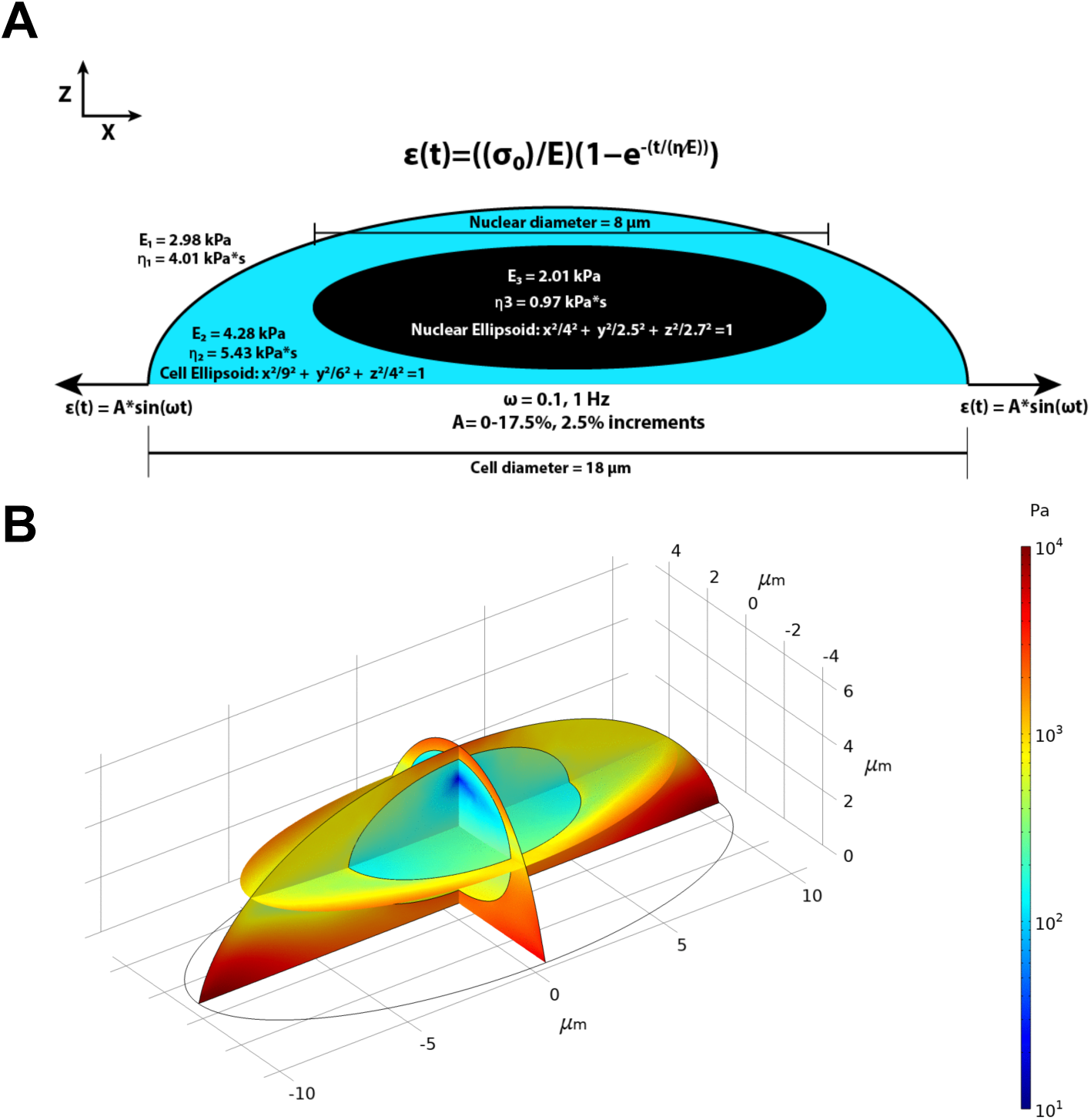
**(A)** Representative diagram of a mesenchymal stem cell with geometric and mechanical parameters for the computational model. Simulated mechanical waveforms include a Sine and Brachial wave at 0.1 Hz or 1 Hz. Maximal strain amplitudes ranged from 0-17.5% in 2.5% increments, and 10 cycles were simulated for each respective waveform. **(B)** Selected three-dimensional diagram of the cell under stress from the COMSOL simulation. Diagram shows the theoretical Von Mises stress in various regions of the cell during the final cycle of a brachial 1 Hz 17.5% maximal strain amplitude mechanical loading regime.

### Computational Simulation and Analysis

The simulation of cell deformation was performed using COMSOL Multiphysics (Comsol, Inc., Burlington, MA; Version 6.0). The components of the cell were modeled as viscoelastic materials with the stress-strain relationship was defined as follows:

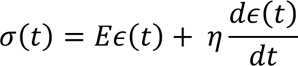

Where σ(t) is the stress, ɛ(t) is the strain, *η* is the viscosity, and *E* is the elastic modulus.^27^ The model used a solid mechanics model for the cytoplasm and nucleus and a membrane model for the cell membrane. Cell model geometry was meshed using the parameters described in **Fig. 1A**. First, the entire cell was meshed with a maximum element size of 0.99 µm and a maximum element growth rate of 1.4. The mesh near the edges of the cell compartments were refined to achieve a higher resolution, with a maximum element size of 0.159 µm **(Supplementary Fig. 1C-F)**.

To apply strain to the cell, biaxial stress consistent with experimental data was applied to the membrane substrate, and the bottom surface of the cell was assumed to be fixed to the membrane, perfectly coupled with the flexible membrane strain, and constrained in the -Z dimension:

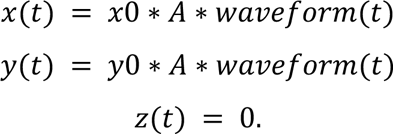

Where x0, y0 are the initial coordinates of any point on the bottom of the cell, A is biaxial strain amplitude and waveform(t) is the applied waveform function.

Time-dependent solutions were generated by using solver settings for output time steps of 0:T/50:10*T. The Transient Generalized alpha method was used for time-stepping with an upper limit on the time step size equal to 1/50 of the period T. The iterative Newton solver with a constant damping factor was applied. Simulation iterations were terminated when the estimated solution-based relative error is less than the specified relative tolerance of 0.001. PARDISO (Parallel Direct Sparse Solver) was used as the direct linear solver used for solving sparse linear systems of equations. The Function Sweep and Parametric Sweep were embedded into the main COMSOL modeling file. Function Sweep was used as the top-level for-loop, computing the solutions for Sine and Brachial waveforms. Parametric sweep was used as the nested for-loop, computing the solutions for all combinations of mechanical loading frequency and maximal strain amplitude.

After generating datasets containing the mechanical response of the model to the application of 10 cycles of mechanical loading with specific waveform, frequency of loading, and maximal strain amplitude, data were sorted to enable comparison between stretch conditions. Data were organized in Microsoft Excel to analyze the resultant mechanical response for each waveform condition. Data columns for each mechanical loading conditions across 10 cycles included: Average von mises stress in the nucleus, estimated strain energy in the nucleus, average true X strain in the nucleus, average true Y strain in the nucleus, average true Z strain in the nucleus, average von mises stress on the nuclear surface, average X strain on the nuclear surface, average Y strain on the nuclear surface, and average Z strain on the nuclear surface. After sorting, data were analyzed to extract maximum, minimum, and average strain values across the 10 cycles of loading for each mechanical loading condition. The effect of each cycle was compared by analyzing the max, min, and average strain values between each cycle of loading. Data plots were generated in Graphpad Prism (GraphPad Software, Inc.), and animation snapshot frames and GIFs were prepared with EZGIF (www.ezgif.com).

### Kelvin-Voigt Model of chromatin deformation

To prepare the model of viscoelastic chromosomes, we applied the Kelvin-Voigt equation and derived it to solve for strain based on an input of stress over time:

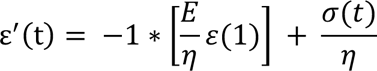

where *ε* is strain, *E* is the elastic modulus of the chromosome, *η* is the viscous damping constant, and *σ* is the input stress.^28^ Following previously described methods, we calculated *E* = 3.82 Pa and *η* **=** 68.81 N•s/m for interphase chromosomes, and *E* = 611.63 Pa and *η* **=** 733.95 N•s/m for mitotic chromosomes.^29^ To calculate the input stress, we assumed perfect coupling and applied the elastic modulus of the nucleus in our model, 2010 Pa, the simulated average X strain in the nucleus values in Pa, to solve *σ **=** ε *E* for each waveform condition across 10 cycles of simulated loading. The model also assumes that all input stress is delivered directly to a chromosome in interphase or mitosis. The spline fit function in MATLAB was used to form continuous functions representing the stress input across 10 cycles of loading for each mechanical waveform condition. Next, the differential equation for the Kelvin-Voigt equation was solved in MATLAB using, yielding axial strain in the X dimension across chromosomes in interphase and mitosis.

## Results

### Preparation of computational model

First, a representative diagram of an individual mesenchymal stem cell was prepared (**Fig. 1A**). Mechanical properties for the cell membrane, cytoplasm, and nucleus were approximated based on a previous study.^27^ Additional literature was referenced to approximate the size and geometry of the cell and nucleus.^24–26^ We included three key regions of the cell, the cell membrane, cytoplasm, and the nucleus, each with their own distinct geometry, elastic modulus, and viscosity. The MSC mechanical parameters were input to COMSOL Multiphysics for the conduction of simulations (**Fig. 1B**). We applied brachial (**Fig. 2A, B**) and sine (**Fig. 2C, D**) waveforms to the model at frequencies of 0.1 and 1 Hz and amplitudes ranging from 0-17.5% maximal strain, in increments of 2.5%. This method generated a large state space of data, including Von Mises stress and strain in the X, Y, and Z dimensions across all possible waveform conditions.

**Figure 2.**
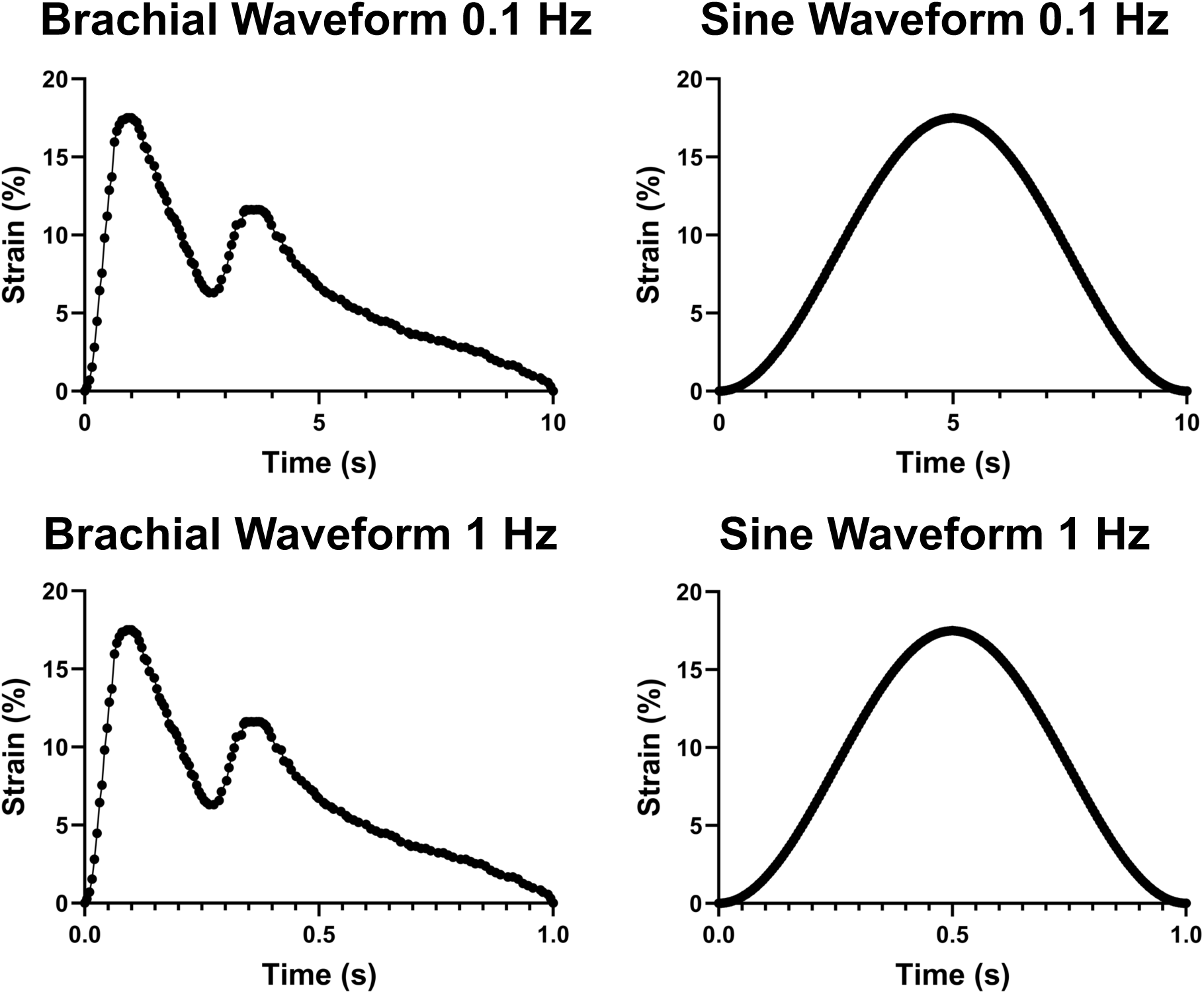
The strain profile of mechanical waveforms simulated in the computational model. The brachial waveform is a waveform that mimics the stretch of the brachial artery during the cardiac cycle (brachial loading) shown at 0.1 Hz or 1 Hz frequency of loading. The sinusoidal waveform is the most commonly used strain waveform in experimental studies (shown at 0.1 Hz or 1 Hz frequency of loading).

### The application of lower frequency mechanical waveforms causes a uniaxial distribution of mechanical strain to the cell nucleus

First, we observed the distribution of strain in the X and Y dimensions across the entirety of the cell. By placing cross-sectional points at the center of the cell diagram, we were able to measure the distribution of mechanical strain across the nucleus during the 10-period cycle of each waveform. We measured the maximal strain delivered to the nucleus during the waveform cycle, for each of the waveform amplitudes at 0.1 and 1 Hz frequency of loading. We found that while maximal strain delivered to the nucleus in the X dimension was similar between 0.1 and 1 Hz waveforms (**Fig. 3A**), maximal nuclear strain in the Y dimension was much greater from 1 Hz waveforms loading compared to those at 0.1 Hz frequency of loading (**Fig. 3B**). We compared the ratio of maximal nuclear strain in the X and Y dimensions, finding that the 1 Hz waveforms generated almost perfectly biaxial nuclear strain, while 0.1 Hz waveforms were strongly uniaxial and biased towards the X dimension of the nucleus, with a max X/Y strain ratio of ∼3.5 (**Fig. 3C; Fig. S1**). This effect was even more strongly pronounced in sine waveforms, with 1 Hz waveforms demonstrating a max X/Y strain ratio of ∼2.5, and 0.1 Hz a max X/Y strain ratio of ∼11, across all simulated amplitudes of strain (**Fig. 3D-F**; **Fig. S2**).

**Figure 3.**
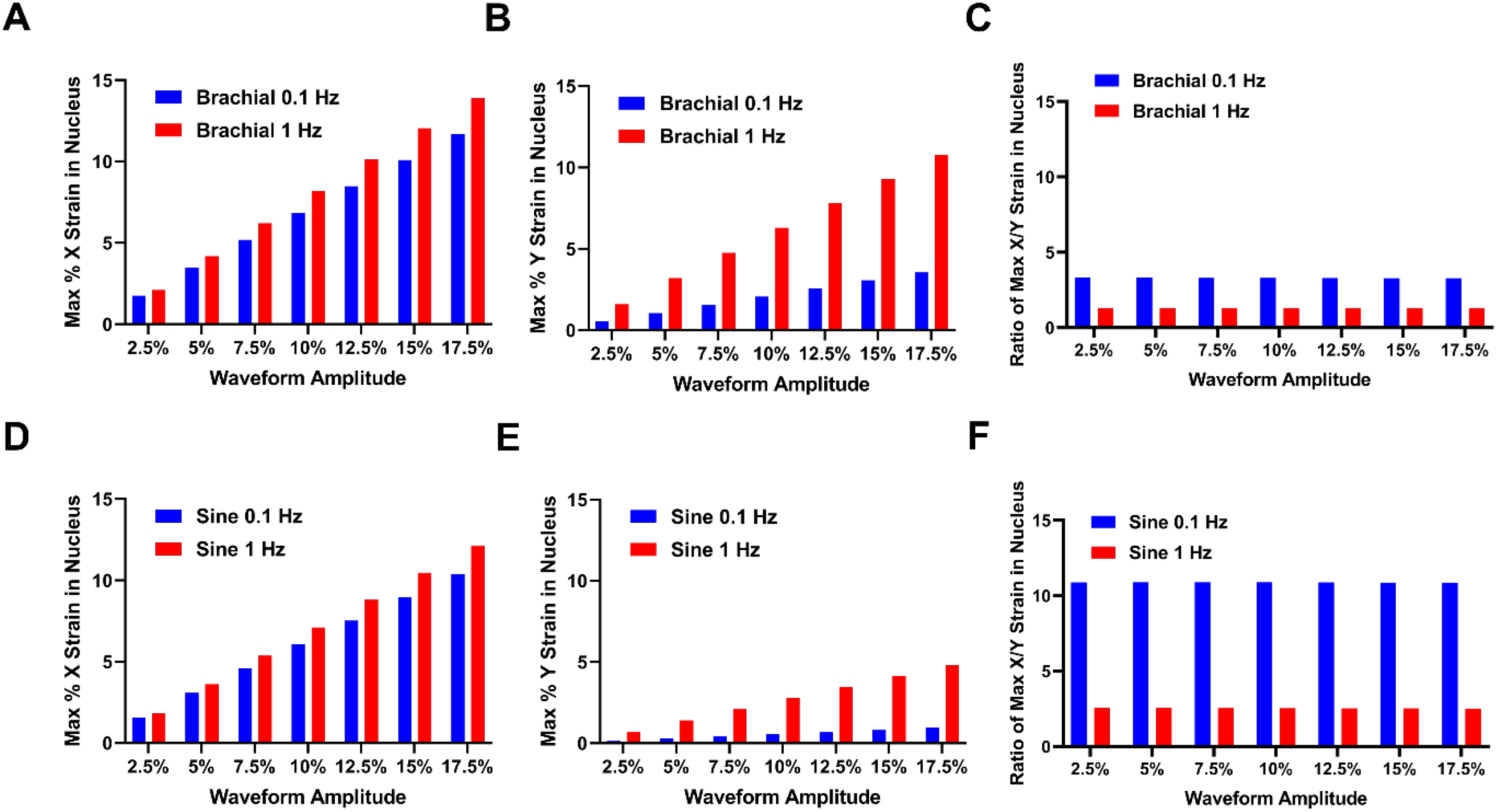
The application of 0.1 Hz mechanical strain causes a uniaxial strain distribution in the cell nucleus. **(A, B)** Maximal nuclear strain for brachial waveforms in the **(A)** X dimension and **(B)** Y dimension. **(C)** The ratio of the maximal X/Y nuclear strain for brachial waveforms. **(D, E)** Maximal nuclear strain for sinusoidal waveforms in the **(D)** X dimension and **(E)** Y dimension. **(F)** The ratio of the maximal X/Y nuclear strain for sinusoidal waveforms.

We repeated the simulations, this time focusing on the distribution of strain to the nuclear surface, representing the nuclear envelope of the cell (**Fig. 4**). We observed a similar trend; nuclear surface strain in the X dimension was similar between 0.1 Hz and 1 Hz waveforms (**Fig. 4A**), but significantly different in the Y dimension (**Fig. 4B**). Once again, we observed that the 1 Hz waveforms delivered biaxial strain to the nuclear surface for both sine and brachial waveforms, while the 0.1 Hz waveforms were strongly uniaxial in the X dimension, with a max X/Y strain ratio of ∼3 (**Fig. 4C; Fig. S3**). The same trend persisted for sine waveforms, with 1 Hz waveforms demonstrating a max X/Y strain ratio of ∼2.5, and 0.1 Hz waveforms with a max X/Y strain ratio of ∼9, across all simulated amplitudes of strain (**Fig. 4D-F**; **Fig. S4**).

**Figure 4.**
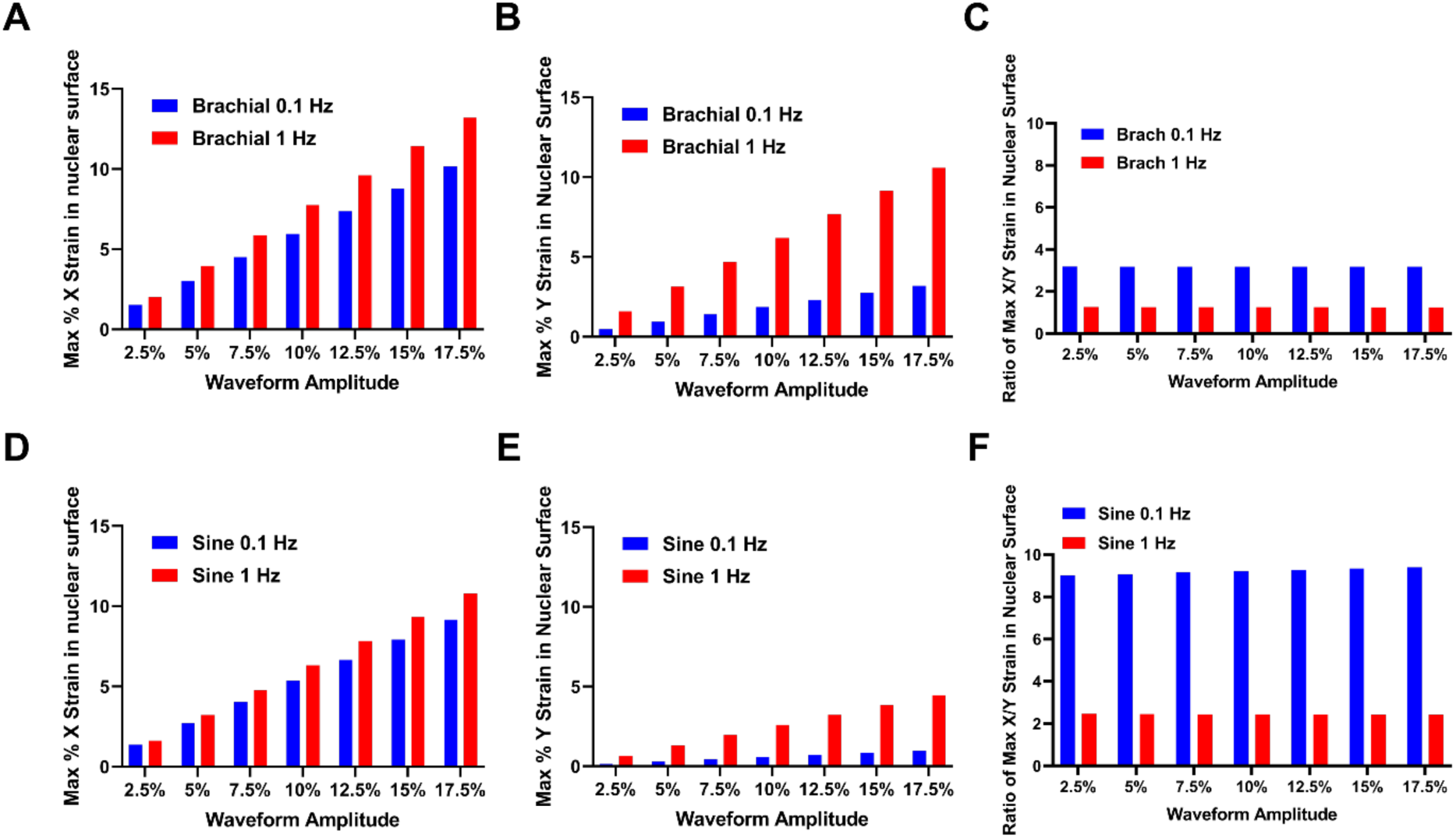
The application of 0.1 Hz mechanical strain causes a uniaxial strain distribution on the nuclear surface. **(A, B)** Maximal nuclear surface strain for brachial waveforms in the **(A)** X dimension and **(B)** Y dimension. **(C)** The ratio of the maximal X/Y nuclear surface strain for brachial waveforms. **(D, E)** Maximal nuclear surface strain for sinusoidal waveforms in the **(D)** X dimension and **(E)** Y dimension. **(F)** The ratio of the maximal X/Y nuclear surface strain for sinusoidal waveforms.

### The nuclear strain profile caused by the first cycle of mechanical loading is significantly different from the strain caused by the remaining cycles of loading

We repeated the simulations of brachial and sinusoidal waveforms at 0.1 and 1 Hz frequency of loading at varying maximal strain amplitudes ranging from 2.5-17.5%. We observed the resultant nuclear strain profile in the X, Y, and Z dimensions, and compared the results across each of the 10 cycles of loading. Results indicate that the strain profile generated by the first cycle of loading differs from nuclear strain generated in cycles 2-10 (**Fig. 5**; **Fig. 6; Fig. S5-S6**). For all mechanical waveforms simulated, the nuclear strain in the X and Y dimensions is initially greater at cycle 1, before decaying in an exponential pattern in subsequent cycles. Conversely, the nuclear strain in the Z dimension demonstrates the opposite effect, with the strain at cycle 1 initially lower, then rapidly increasing to a relatively constant value across cycles 2-10. The differential strain caused by this cycle effect was more pronounced in 1 Hz mechanical waveforms but persisted across the 0.1 Hz waveforms as well. The exponential decaying pattern of the strain across the 10 cycles suggests that the nuclear strain profile reaches a quasi-steady state by cycle 10, which likely persists throughout the remainder of the mechanical conditioning treatment.

**Figure 5.**
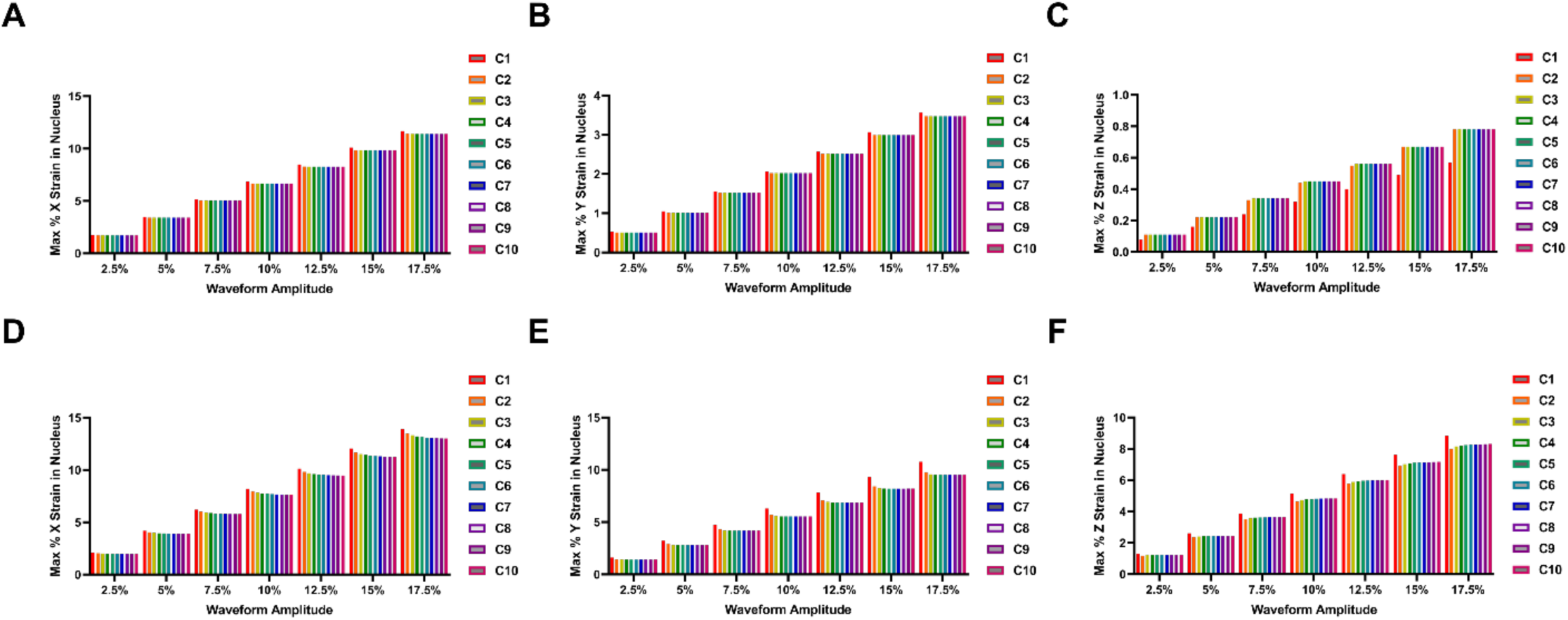
Effect of cyclic brachial loading on mechanical strain in the nucleus. **(A-C)** Simulated strain profile across 10 cycles of the brachial waveform at 0.1 Hz frequency of loading for nuclear strain in the **(A)** X, **(B)** Y, and **(C)** Z dimensions. **(D-F)** Simulated strain profile across 10 cycles of the brachial waveform at 1 Hz frequency of loading for nuclear strain in the **(D)** X, **(E)** Y, and **(F)** Z dimensions.

**Figure 6.**
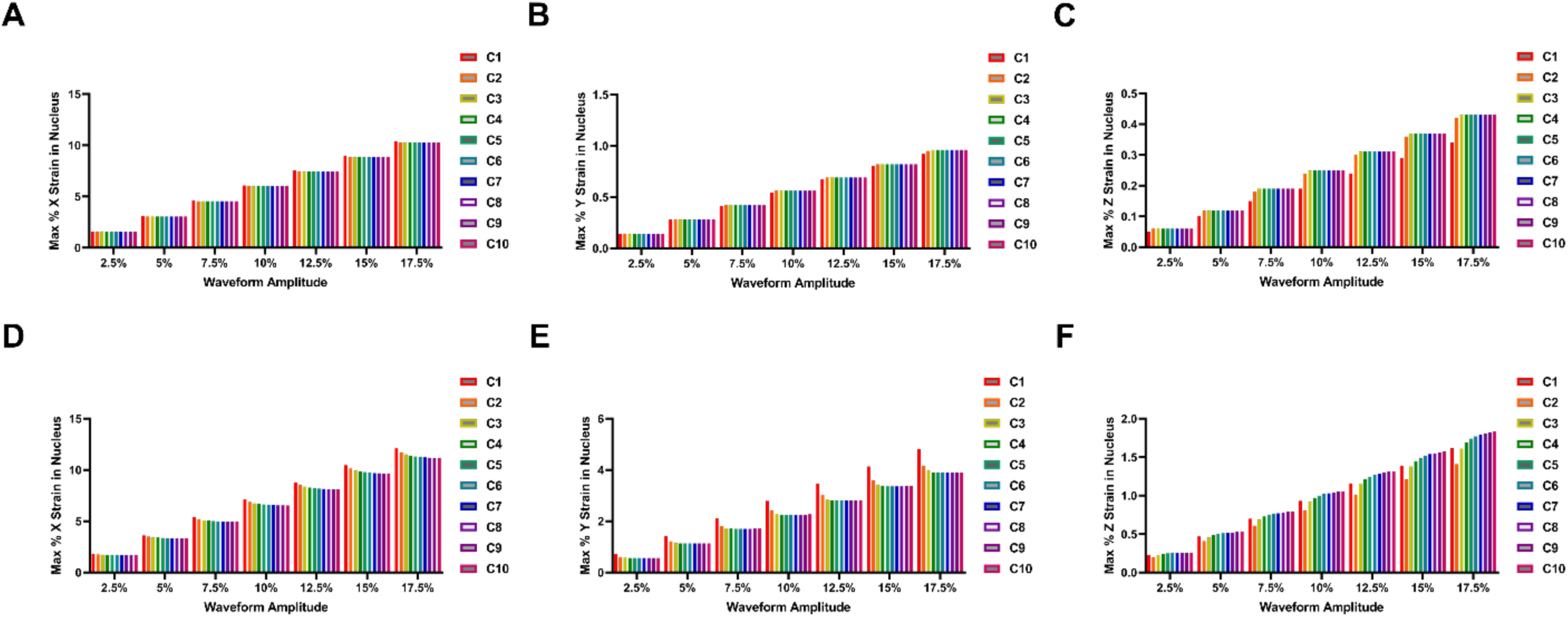
Effect of cyclic sinusoidal loading on mechanical strain in the nucleus. **(A-C)** Simulated strain profile across 10 cycles of the sinusoidal waveform at 0.1 Hz frequency of loading for nuclear strain in the **(A)** X, **(B)** Y, and **(C)** Z dimensions. **(D-F)** Simulated strain profile across 10 cycles of the sinusoidal waveform at 1 Hz frequency of loading for nuclear strain in the **(D)** X, **(E)** Y, and **(F)** Z dimensions.

### The average nuclear strain profile across the final cycle of loading demonstrates multi-dimensional generation of tensile and compressive strain

Next, we analyzed the mean mechanical stress applied to the nucleus in the X, Y, and Z axes throughout the tenth cycle of both sine and brachial 0.1 Hz wave patterns. This was done to gain a deeper insight into why brachial wave patterns appear to trigger more genetic and phenotypic alterations compared to sine wave patterns of comparable magnitude. Cycle 10 was selected for this analysis as we anticipate that the mechanical strain profile during this cycle is representative of steady state conditions that persist throughout the mechanical conditioning treatment. We observed that the X strain followed expected strain patterns generated by the waveform, while Y strain patterns demonstrated both tension and compression during the cycle, and Z strain patterns are almost fully compressive (**Fig. 7**). These results were unexpected and lead us to several hypotheses.

**Figure 7.**
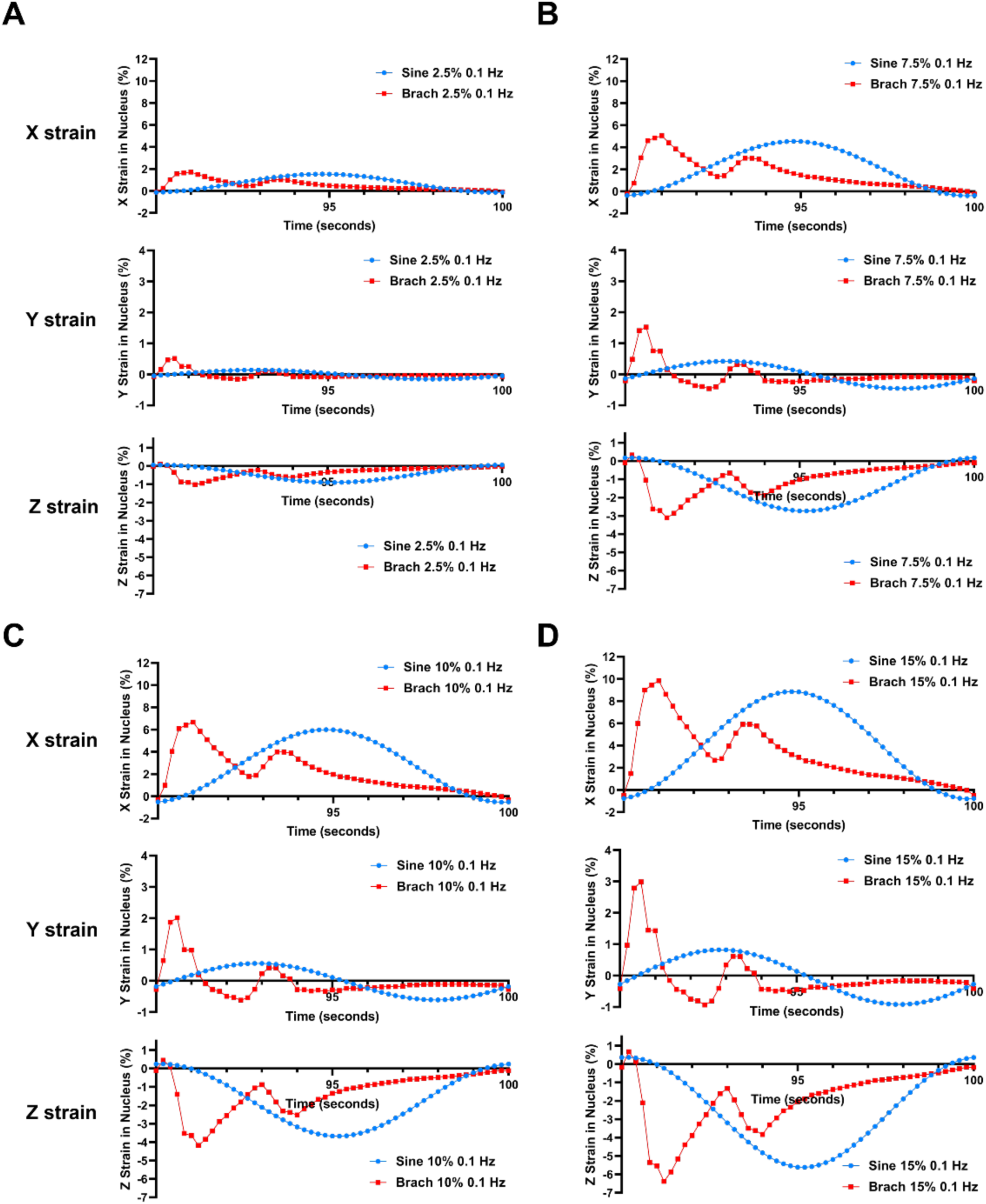
The average nuclear strain profile across the final cycle of loading demonstrates multi-dimensional generation of tensile and compressive strain. The average nuclear strain in the X, Y, and Z dimensions across cycle 10 were analyzed for 0.1 Hz sine and brachial waveforms of **(A)** 2.5% maximal strain amplitude, **(B)** 7.5% maximal strain amplitude, **(C)** 10% maximal strain amplitude, and **(D)** 15% maximal strain amplitude.

First, the results support our previous claim that brachial waveforms generate a more biaxial strain profile than sine waveforms, likely due to the initial rapidly increasing application of strain in the brachial cycle. Sine waveforms, in contrast, deliver strain much more gradually over the course of the cycle, enabling the cell to relax and behave more like a viscous fluid,^30^ demonstrated by their out-of-phase strain response in the Y dimension, represented as the short axis of the cell. All simulated waveforms demonstrated a compressive strain profile in the Z dimension, suggesting that the cell is being flattened due to tensile strain along the X and Y axes, representing the long and short axes of the cell, respectively. In our simulations, low amplitudes of strain (2.5%) caused minimal compressive effects in the Y and Z dimension (**Fig. 7A, Fig. S7**), while mid-range (**Fig. 7B, 7C; Figs. S8-S9**), and high amplitudes (**Fig. 7D; Fig. S10-S12**) generated compressive strain in the Z dimension of similar magnitude to tensile strain in the X dimension. Thus, these data may provide an explanation for why cyclical strain of low magnitudes causes less alterations in mechanotransductive signaling, as demonstrated in our previous study.^23^ The simulated 10 cycles of 1 Hz mechanical loading generated a similar magnitude of strain deformation as the 0.1 Hz waveforms (**Fig. 8**). 1 Hz mechanical loading of interphase chromosomes generated up to 10X total magnitude of deformation after 10 seconds (**Fig. 8A, C**), matching the behavior of the interphase chromosomes under 0.1 Hz loading. Similarly, 1 Hz loading of mitotic chromosomes generated up to ∼17.5% deformation after 10 seconds (**Fig. 8B, D**), matching the behavior of the mitotic chromosomes under 0.1 Hz loading, and effectively following the deformation of the silicone substrate.

**Figure 8.**
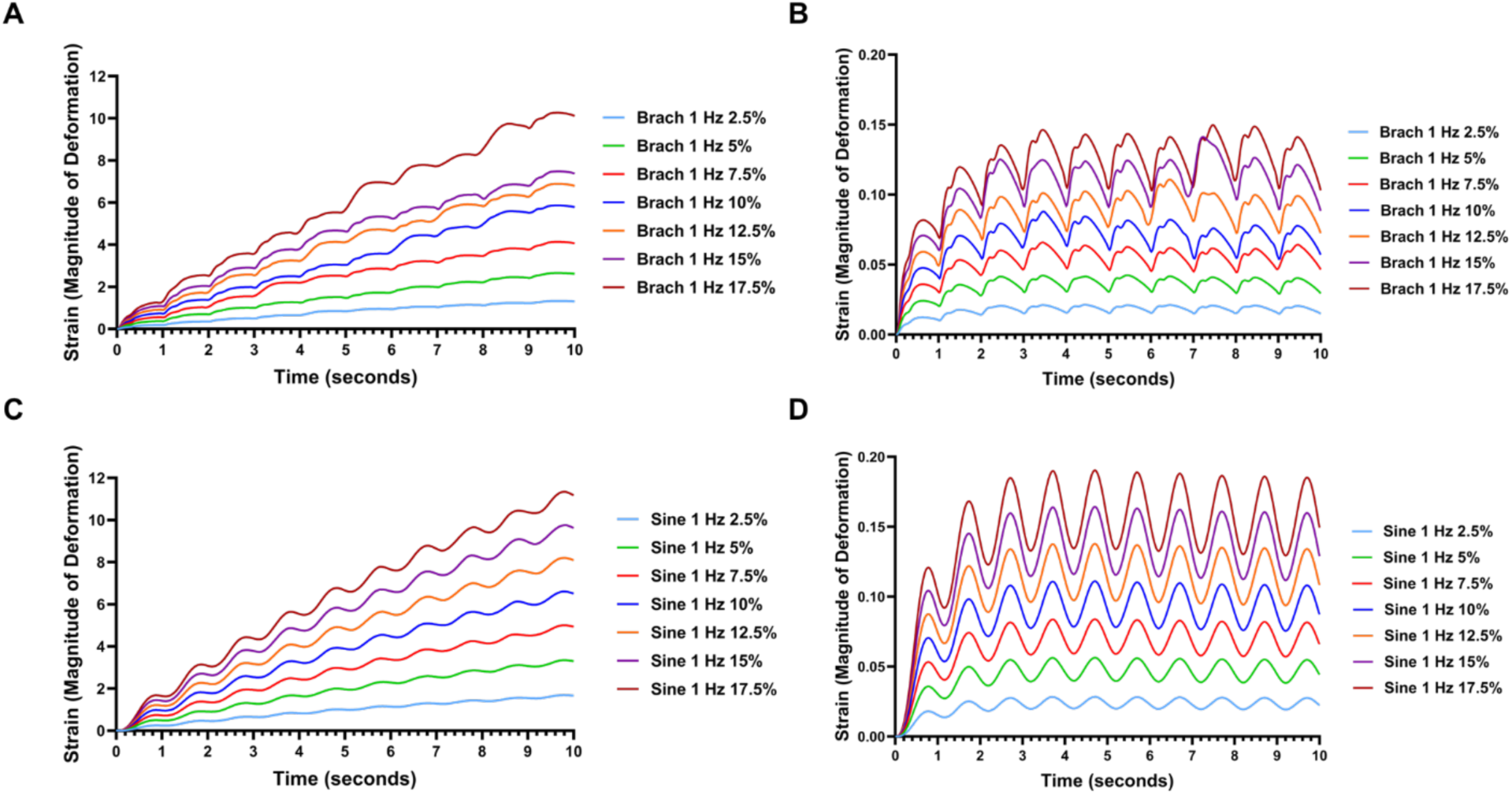
Estimated strain magnitude applied to chromatin in viscoelastic model at 1 Hz frequency of loading. Strain magnitude applied in the X dimension across 10 cycles of mechanical loading at 1 Hz for **(A)** brachial waveform loading of interphase and **(B)** mitosis chromatin. Strain magnitude applied in the X dimension across 10 cycles of mechanical loading at 1 Hz for **(C)** sine waveform loading of interphase and **(D)** mitosis chromatin.

### Midrange strains at 0.1Hz are theoretically sufficient to mechanically unravel chromosomes at interphase without chromatin damage

Next, we incorporated a viscoelastic model of chromosomes under axial stretch to determine their extension under 10 cycles of cyclic mechanical loading at various waveforms, frequencies, and strain amplitudes. Recent work has proposed a viscoelastic model of chromosomes and their mechanical properties during interphase and mitosis.^29,31^ This work used next-generation sequencing techniques and mathematic simulations to characterize the folding and structural conformations of individual chromosomes. Simulations generated force-extension curves demonstrating that mitotic chromosomes behave about 10-fold stiffer than interphase chromosomes.^29^ We followed the results of this study to generate a representative Kelvin-Voigt model of viscoelasticity in MATLAB for evaluating the extension of each type of chromatin during cyclic sine and brachial mechanical loading.^32^ We observed that, within the 10 cycles of 0.1 Hz simulated mechanical loading, interphase chromosomes reached oscillatory steady-state behavior in approximately 50 seconds (**Fig. 9A, C**), while mitotic chromosomes reached steady-state at approximately 3 seconds (**Fig. 9B, D**). It has been reported that chromosomes mechanically unravel when extension reaches roughly double its native length, with damage occurring at approximately three times its native length.^29^ Thus, strains around two-fold the initial length could be ideal for mechanically inducing strain We assumed that after reaching oscillatory steady state the cell adapts to only experience the strains applied each cycle and plotted the strain for each cycle under the various conditions (**Fig. 9E, F**). We found that for the 0.1 Hz sine wave form, strains of 7.5-10% were in this range of 2-3 fold extension. Interestingly, the 0.1 Hz brachial waveform had a broader set of strains in this region, with 7.5%- 12.5% strain leading to nuclear strains around two-folder. For the mitotic stage, none of the strains induced the range of extension for unfolding chromatin. Thus, this optimal level of strain corresponds well to the optimal conditions for inducing vascular differentiation in MSCs.^23^

**Figure 9.**
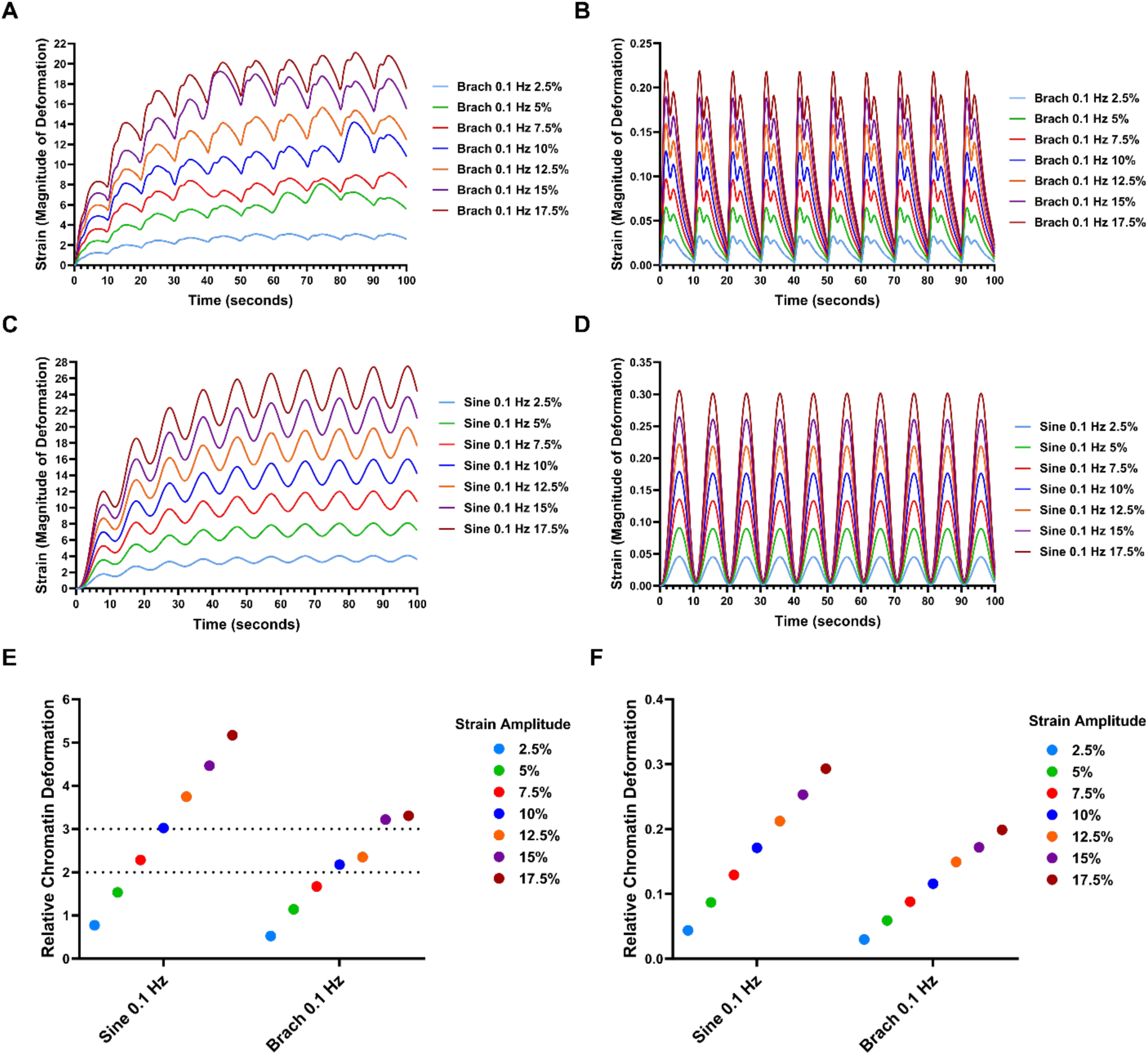
Estimated strain magnitude applied to chromatin in viscoelastic model at 0.1 Hz frequency of loading. Strain magnitude applied in the X dimension across 10 cycles of mechanical loading at 0.1 Hz for **(A)** brachial waveform loading of interphase and **(B)** mitosis chromatin. Strain magnitude applied in the X dimension across 10 cycles of mechanical loading at 0.1 Hz for **(C)** sine waveform loading of interphase and **(D)** mitosis chromatin. **(E)** Relative deformation of interphase chromatin across cycle 10 of simulated sine and brachial waveforms at 0.1 Hz frequency of loading. **(F)** Relative deformation of mitosis chromatin across cycle 10 of simulated sine and brachial waveforms at 0.1 Hz frequency of loading.

## Discussion

Recent advances in genetic sequencing, imaging, and computational techniques have enabled researchers to better understand the complex role of mechanical forces in regulating cell homeostasis. Numerous studies have deeply investigated specific signaling pathways of mechanotransduction and mechanoadaptation, in which cells respond to external mechanical forces through population-wide alignment,^33^ phenotypic and lineage-specific changes,^34^ and transcriptomic and epigenetic modifications.^13–15^ However, there remains a need for thorough investigation of how the specific type of mechanical force applied to the cell differentially changes its response. Emerging literature suggests that a better understanding of the cell’s response to mechanical strain may lead to the engineering of exciting new therapeutic applications,^35,36^ directed differentiation and maintenance of stem cell phenotypes,^37^ and a deeper understanding of the role of mechanical forces in healthy tissue homeostasis and disease progression.^38,39^ In this work, we created a computational model of the application of mechanical strain to a single mesenchymal stem cell (MSC) cultured on a flexible silicone membrane. To consider the differential mechanical response caused by different types of mechanical forces, we performed simulations involving a broad state space of mechanical strain amplitudes, waveform patterns, and frequencies of loading. We investigated the mechanical deformation of the cell membrane, cytoplasm, and nucleus across all three dimensions of the cell. This study provides a detailed characterization of a cell’s mechanical response to physiological forces and proposes mathematical reasoning for why specific types of forces generate more significant biological alterations in mechanotransduction and gene expression.

Cell attempts to model a cell’s mechanical properties and response to forces are largely focused on nuclear and chromatin mechanics. These studies vary substantially in the parametrization of a cell’s mechanical properties, including discrete and mixed-phase models of cellular compartments, porous solid models of the filamentous network structure of the cytoskeleton and LINC complex, and viscoelastic models of the force and deformation of chromatin in various states.^40,41^ Several computational studies consider detailed mechanical sub-sections of the nucleus and their interaction during mechanotransductive signaling, including the nuclear envelope, nucleoli, and chromatin strands of various stiffness and compaction.^16,17,42^ These works provide key detail regarding the importance of biological stimuli like histone modifications in determining the nucleus’ response to mechanical strain. Several studies have used image-derived modeling techniques to better understand the heterogeneous mechanical behavior of the nucleus, dependent on the assortment of low-stiffness euchromatin and high-stiffness heterochromatin.^13,14^ These studies characterized the nuclear response to shear strain and its relationship with mechanotransduction, but did not consider other types of mechanical forces, such as tensile and compressive strain, or physiological patterns of force, that are commonly present *in vivo*. Other recent work has used worm-like-chain models of DNA and computational tools to investigate the deformation of chromatin under mechanical forces and the resultant changes in histone binding complexes and chromosome-wide reorganization.^17,29^ These works provided an excellent reference point for this study, enabling a deeper analysis into how different amplitudes and patterns of forces may result in altered nuclear re-organization. In this regard, our work provides a framework for understanding how applied dynamic mechanical strains to cells on flexible substrates is transferred to the nucleus.

A major goal of these studies was to understand how the differences in mechanical loading parameters alter the mechanical forces on the nucleus that may lead to chromatin remodeling. We compared simulation of 0.1 Hz and 1Hz mechanical loading waveforms, and found that low frequency waveforms (0.1 Hz) resulted in a more uniaxial distribution of strain in the nucleus compared to higher frequency waveforms (1 Hz) that had a more biaxial distribution of strain in the nucleus. Previous studies have demonstrated that cells tolerate uniaxial strain better than biaxial strain, as biaxial strain at high magnitudes may cause permanent cytoskeletal deformation and destruction of key mechanotransductive signaling elements such as the LINC complex.^43–45^ Furthermore, uniaxial strain is known to induce cellular alignment, while biaxial strain does not cause this effect.^46^ By performing the simulations over 10 cycles for each waveform, we also observed that the initial cycles caused a differentially applied mechanical strain profile compared to subsequent cycles, with strain profiles gradually reaching a steady state by cycle 10. By mapping the average strain in the nucleus over the 10^th^ cycle, we observed that all simulated waveforms, in particular brachial patterns with mid-high (7.5-17.5%) amplitudes of strain, caused multi-dimensional tensile and compressive effects in the X, Y, and Z dimensions of the cell. Recent studies have shown that compressive stresses to cell nuclei induce lasting changes in gene expression through several mechanotransductive mechanisms.^42^ Compressive forces applied to cell nuclei were found to deform the nuclear envelope and reduce its mechanical restriction, upregulating nuclear transport of ATR (Ataxia telangiecstasia and Rad3 related) and Yap/Taz, influencing the regulation of chromatin condensation/decondensation and protecting chromatin integrity.^47,48^ Thus, these data may provide an explanation for why cyclical strain of low magnitudes causes less activation of mechanotransductive signaling.^23^

Using a Kelvin-Voigt viscoelastic model of the chromatin, we estimated the magnitude of extension delivered to individual chromosomes at various mechanical loading conditions. The mitotic chromosomes behaved like stiff solids, and essentially followed the displacement of the silicone membrane substrate. However, interphase chromosomes were shown to extend by several orders of magnitude, with strain extension rapidly increasing after each subsequent cycle of loading before reaching steady-state-like behavior at cycle 5 (∼50 seconds). Additionally, hysteresis of the interphase chromosomes was observed at later loading cycles, leading to slight deviations from the expected steady-state behavior and generating differential stress-strain responses. This residual memory of deformation phenomena has been previously reported in chromatin under mechanical strain, and may be biologically significant during chromatin reorganization processes related to cell migration, differentiation, aging, and disease.^49^ These results further demonstrate that the application of repeated cyclic loading, rather than a one-time linearly increasing application of strain, is key in developing mechanical conditioning regimes that cause directed reorganization of chromatin.

Furthermore, this study revealed that the strain amplitude to mechanically unravel chromosomes requires at least 7.5% maximal strain in both sine and brachial loading conditions. Recent work reported that chromosomes unravel when their extension reaches roughly double their native length.^29^ This process is known to be reversible, and relaxation of force causes dissociated nucleosomes to reassemble and reset the organization of the chromatin.^17,50^ Therefore, we hypothesize that specific loading regimes may reset chromatin organization by extending chromatin strands, removing nucleosomes, and enabling re-organization during periods of mechanical relaxation. Waveforms with strain amplitudes < 7.5% in our study are shown to cause oscillatory extension of less than 1.5x the length of the chromosome at the beginning of each loading cycle. Therefore, these low-end strain waveforms may not be extending chromosomes sufficiently to cause nuclear reorganization and sustained transcriptional alterations. Higher strain amplitude waveforms (>15%) demonstrate 3-4x extension of chromosome length within each loading cycle, which may damage crucial mechanosensitive elements, such as the cytoskeleton and LINC complex.^43–45^ Mid-range strain amplitude waveforms (7.5% - 12.5%) therefore may cause nuclear deformation sufficient for mechanically unraveling chromosomes, with lower risk of causing permanent damage and apoptosis. This observation may explain why waveforms with middle ranges of maximal strain cause the most significant benefits in MSC regenerative properties and nuclear Yap/Taz signaling, as reported in our previous study.^23^

In order to create the model, we made several important assumption and simplifications that are limitations. The model that we employed is highly specific in that it utilizes geometric and mechanical properties of a single type of cell (MSC), grown on a perfectly flat flexible silicone membrane. These properties likely vary between cells, based on cell state and adapt to mechanical loading over the long term. Additionally, this study did not include detailed sub-compartments of the nucleus, such as the nuclear lamina and LINC complex. Studies have shown that the lamina function as shock absorbers for the nucleus,^51^ and models that consider the nucleus as a biopolymeric shell may more accurately replicate nuclear elasticity and response to strain.^52^ Furthermore, while our study maintains fixed viscoelastic properties, recent work has demonstrated that extracellular mechanical forces result in the repositioning and rheological adaptation of chromatin, causing dynamic changes in nuclear viscoelasticity.^12,53,54^ In addition, we made significant simplifications about how force was transferred to chromatin. In true physiological conditions, there lists exists dynamic coupling and de-coupling interactions between the various mechanical compartments of the cell and the extracellular matrix, these interactions are likely adaptive to the levels of strains and remodel the cell structure over the longer term.^55^

Despite the limitations, the results of this study may be useful in understanding the mechanical forces that are present in the cell under stretching on a flexible membrane. In terms of our experimental findings, this work provides rationale as to why the brachial wave form at 0.1 Hz with maximal strains around 7.5% on the cells through mechanical stretch may be optimal for causing effects on chromatin within the nucleus.^23^ The results of this study provide a guide for future cell stretching experiments and therapeutic strategies, demonstrating the importance of selecting specific mechanical parameters in developing mechanical conditioning regimes and informing models of nuclear mechanotransduction. These efforts may generate new methods for engineering patient-specific mechanobiological treatments for enhancing the efficacy of cell-based therapeutics.

## Supporting information

Supplemental Info Part 1

## Acknowledgements

The authors gratefully acknowledge funding through the DOD CDMRP (W81XWH-16-1-0580; W81XWH-16-1-0582) and the National Institutes of Health (1R21EB023551-01; 1R21EB024147-01A1; 1R01HL141761-01) to ABB.

## Author Contributions

MWM and ABB designed research, performed research, analyzed data and wrote the paper. DA performed research, analyzed data and wrote the paper. AD and AS performed research and analyzed data.

## Supplemental Information

The supplemental information for this manuscript can be downloaded at this link: https://drive.google.com/drive/folders/1NdlHkBdclVvusKXAoCeus3prVdnjKnkj?usp=sharing

