## Supplemental Info Part 1 for "A Computational Model of Mechanical Stretching of Cultured Cells on a Flexible Membrane"

### Slide 1
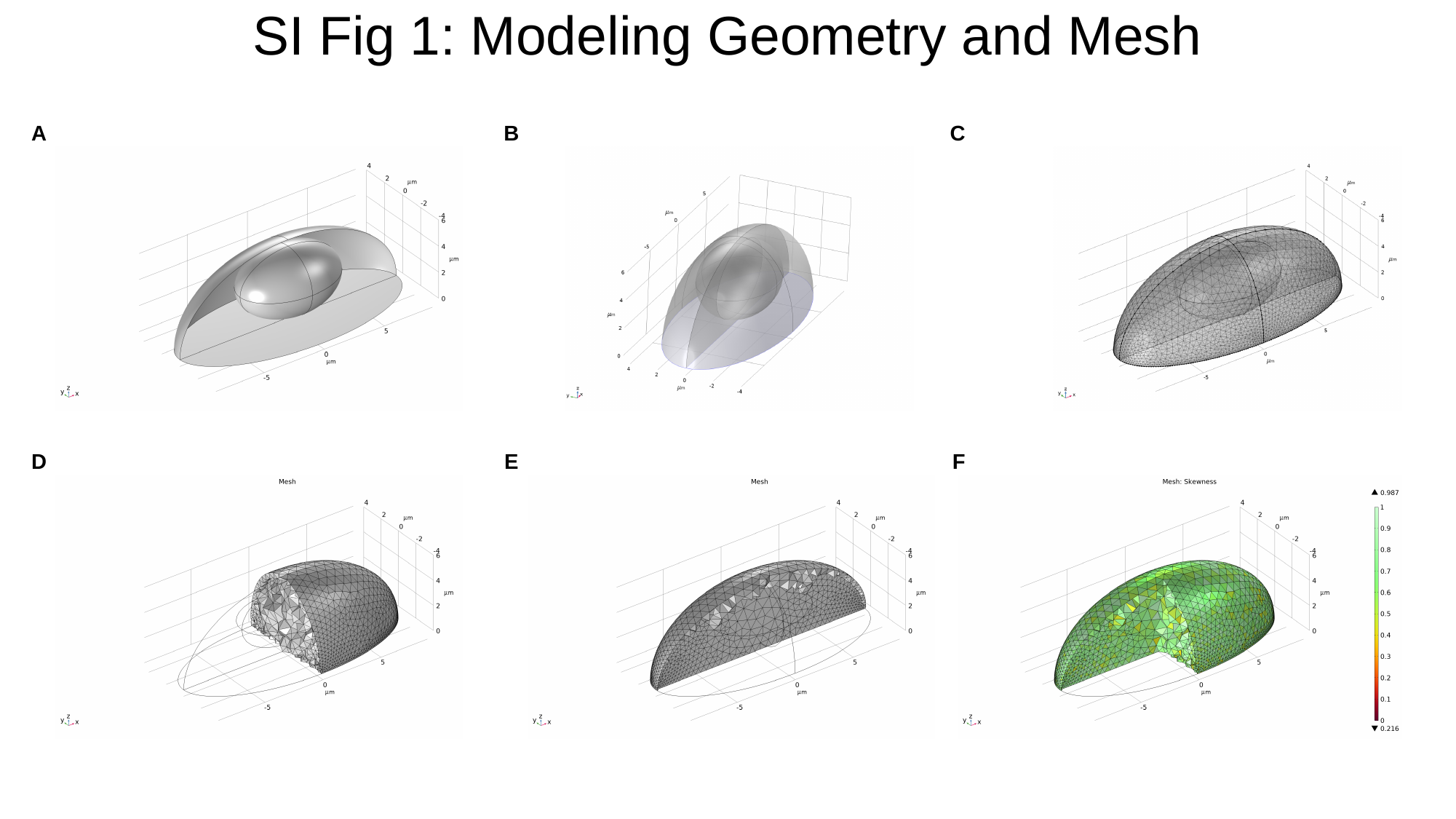

SI Fig 1: Modeling Geometry and Mesh
C
B
A
E
F
D

### Slide 2
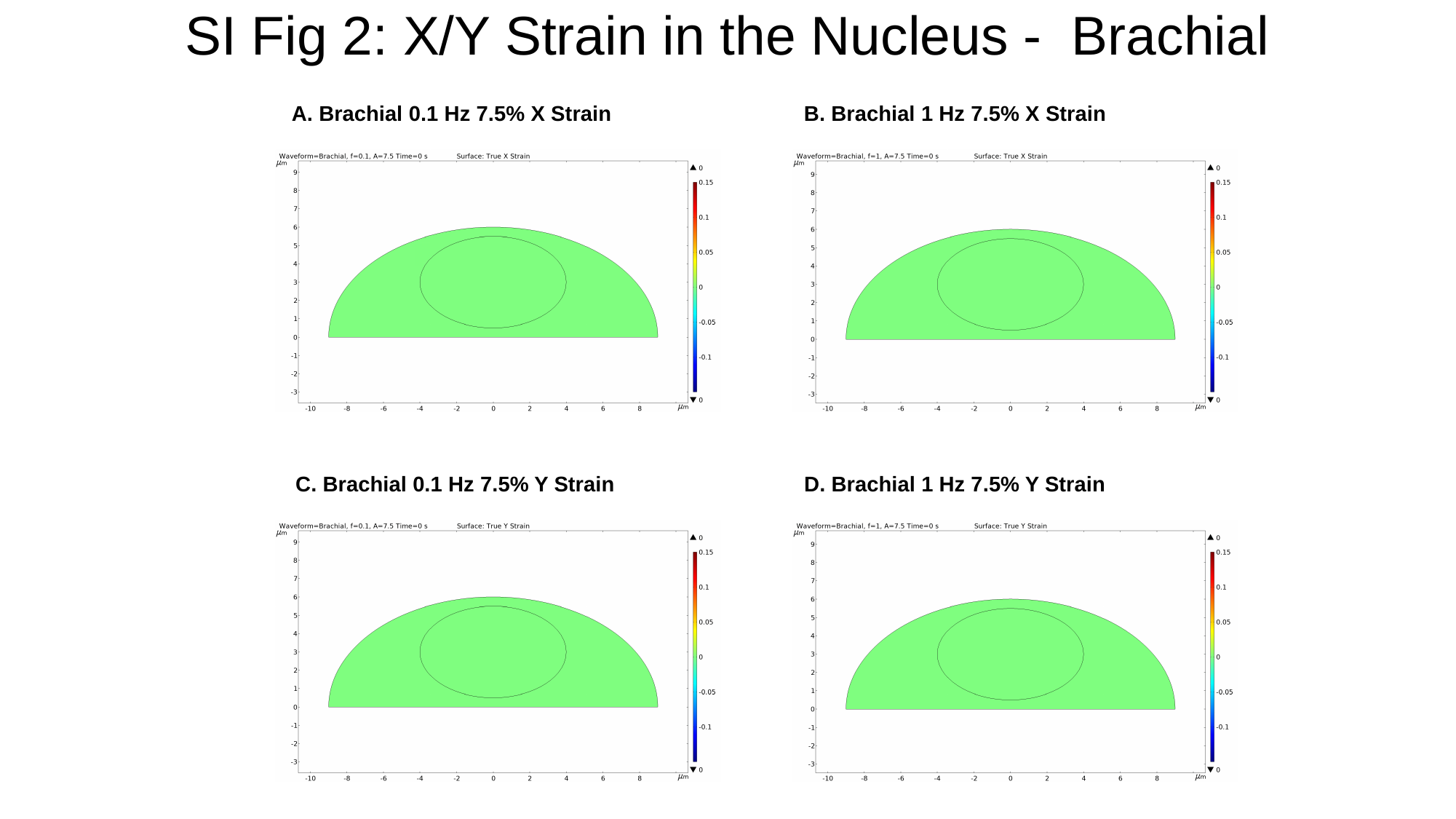

SI Fig 2: X/Y Strain in the Nucleus - Brachial
A. Brachial 0.1 Hz 7.5% X Strain
B. Brachial 1 Hz 7.5% X Strain
C. Brachial 0.1 Hz 7.5% Y Strain
D. Brachial 1 Hz 7.5% Y Strain

### Slide 3
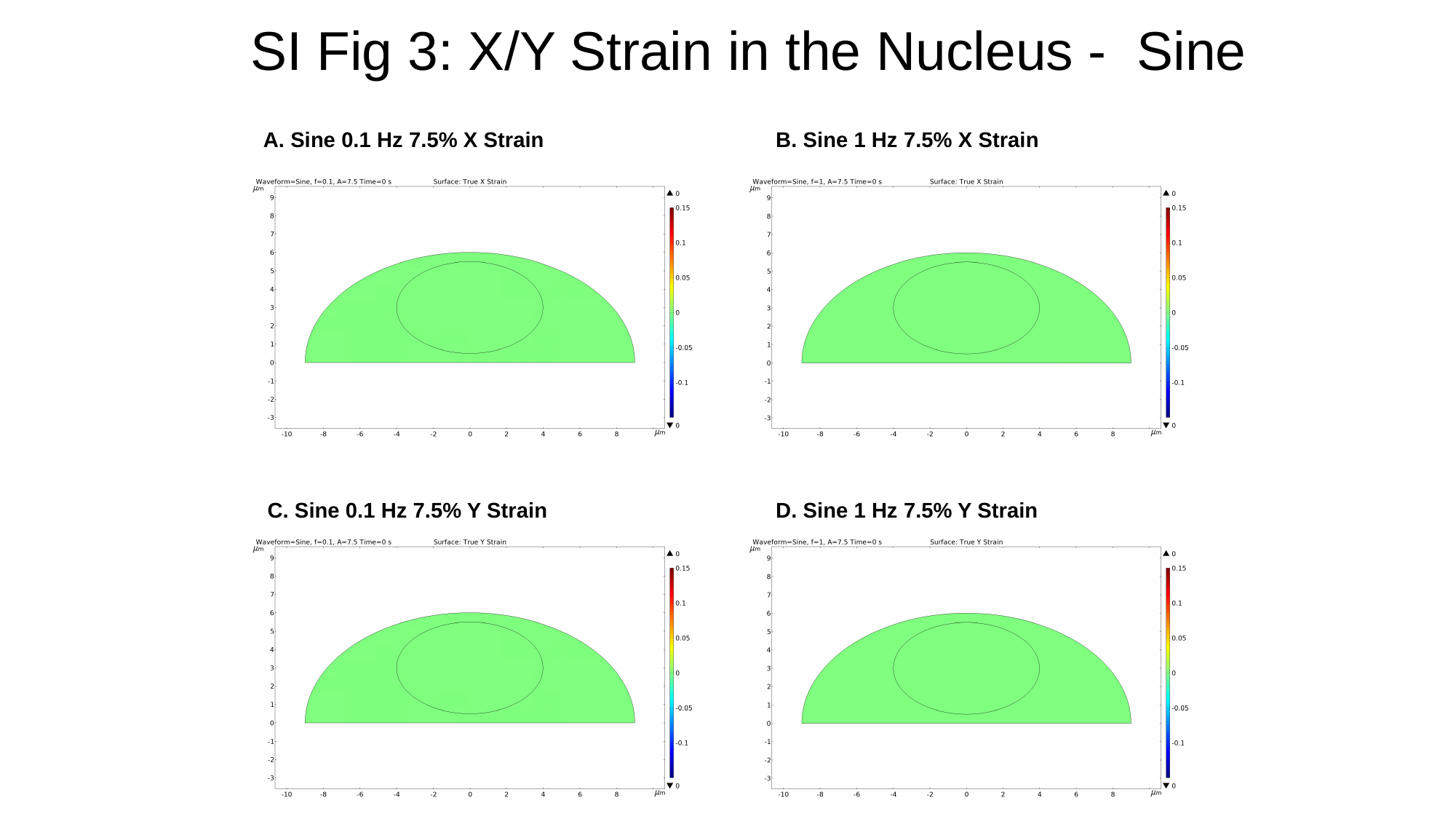

SI Fig 3: X/Y Strain in the Nucleus - Sine
A. Sine 0.1 Hz 7.5% X Strain
B. Sine 1 Hz 7.5% X Strain
C. Sine 0.1 Hz 7.5% Y Strain
D. Sine 1 Hz 7.5% Y Strain

### Slide 4
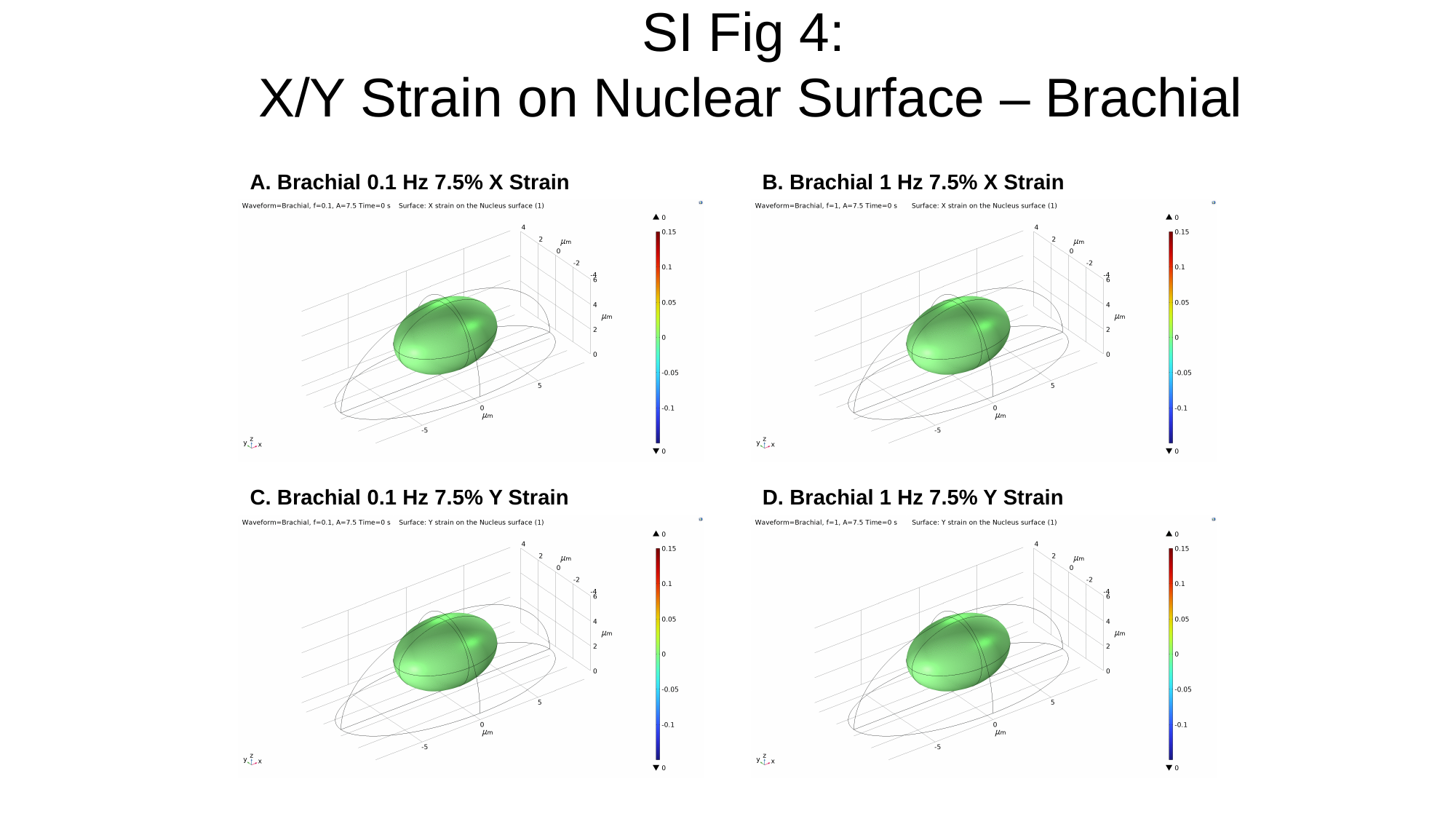

SI Fig 4:
X/Y Strain on Nuclear Surface – Brachial
A. Brachial 0.1 Hz 7.5% X Strain
B. Brachial 1 Hz 7.5% X Strain
D. Brachial 1 Hz 7.5% Y Strain
C. Brachial 0.1 Hz 7.5% Y Strain

### Slide 5
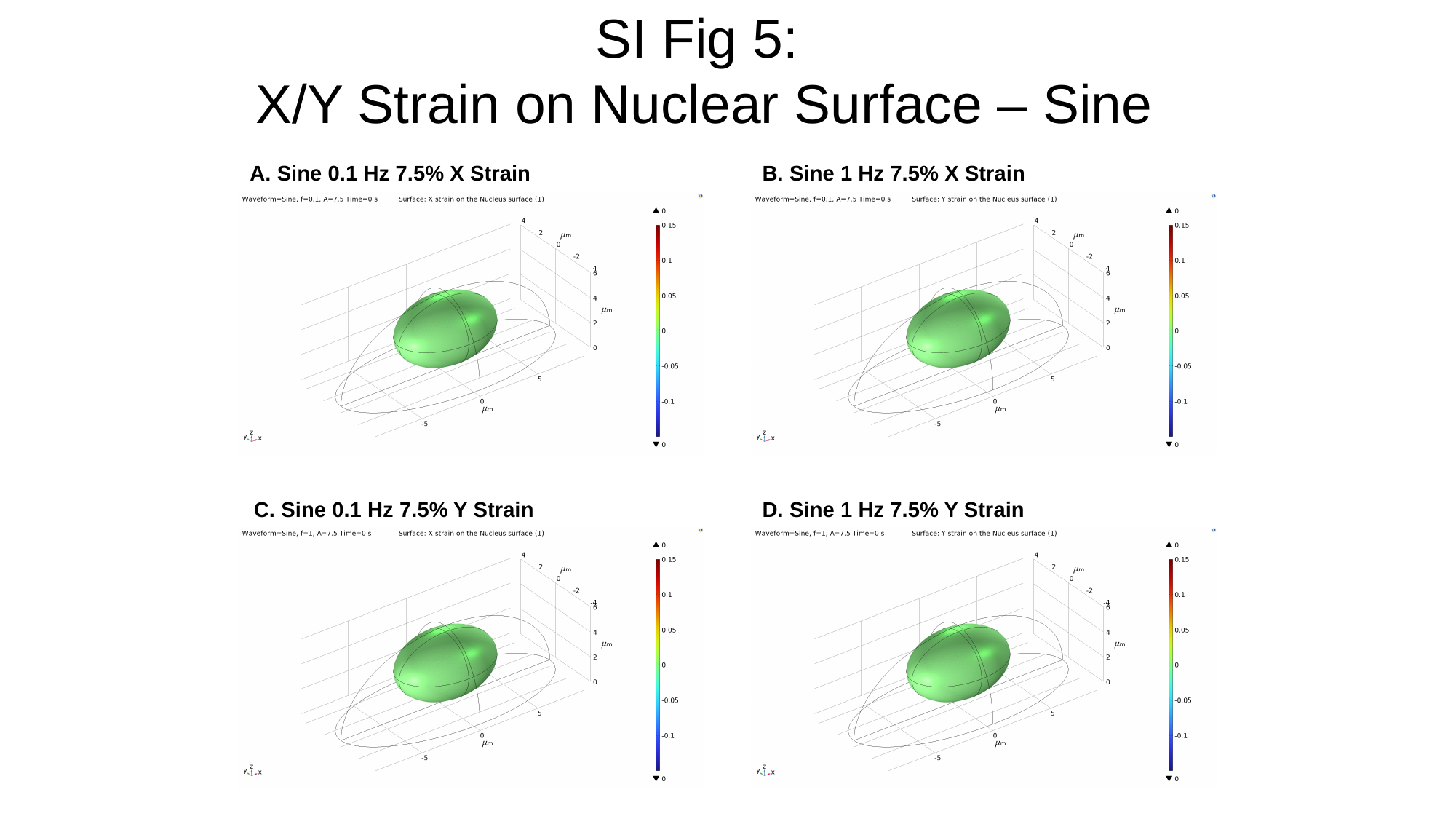

SI Fig 5:
X/Y Strain on Nuclear Surface – Sine
B. Sine 1 Hz 7.5% X Strain
A. Sine 0.1 Hz 7.5% X Strain
C. Sine 0.1 Hz 7.5% Y Strain
D. Sine 1 Hz 7.5% Y Strain

### Slide 6
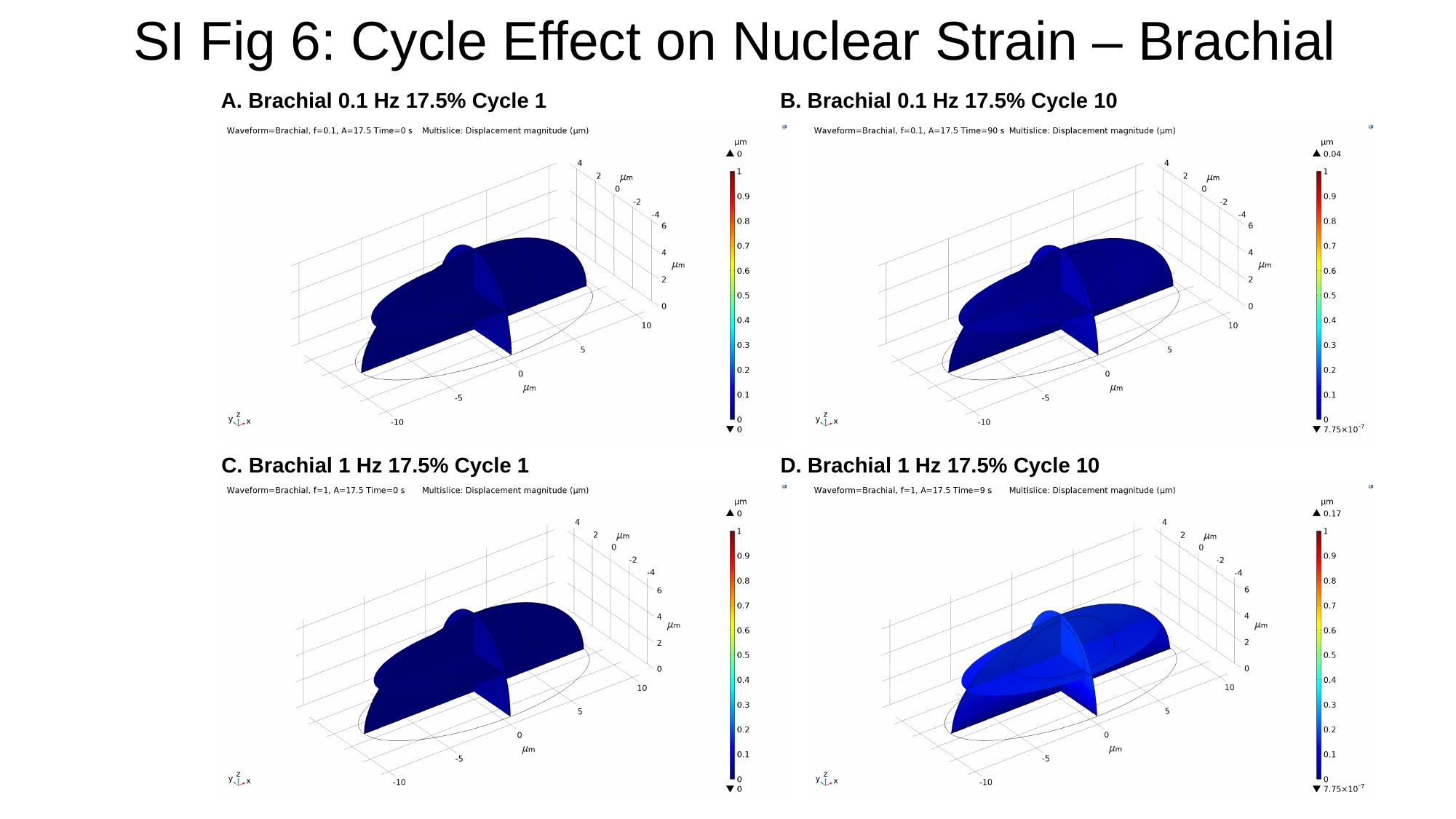

SI Fig 6: Cycle Effect on Nuclear Strain – Brachial
B. Brachial 0.1 Hz 17.5% Cycle 10
A. Brachial 0.1 Hz 17.5% Cycle 1
C. Brachial 1 Hz 17.5% Cycle 1
D. Brachial 1 Hz 17.5% Cycle 10
